# Mitochondrial antigen-specific CD8⁺ T cells drive dopamine neuron neurodegeneration

**DOI:** 10.1101/2024.02.26.582098

**Authors:** Moustafa N Elemeery, Alex Tchung, Salix Boulet S, Nicolas Giguère, Sarah Mezrag, Jean-Francois Daudelin, Amandine Even, Abigail Ralph, Sriparna Mukherjee, Claudie Beaulieu, Diana Matheoud, Jo-Ann Stratton, Nathalie Labrecque, Louis-Eric Trudeau

## Abstract

The progressive degeneration of dopamine (DA) neurons drives motor symptoms in Parkinson’s disease (PD). Whether this neuronal degeneration is due to cell-autonomous dysfunctions in DA neurons or to death signals generated by other cell types is a key problem to address. Recent evidence suggests that loss of function of the protein PINK1, linked to early-onset forms of PD, enhances the presentation of self-derived mitochondrial antigens, which induces the response of autoreactive CD8^+^ T cells. Whether mitochondrial antigen-specific CD8^+^ T cells alone are sufficient to induce nigrostriatal dysfunction has not been directly tested. Here we performed adoptive transfer of mitochondrial antigen-specific CD8^+^ T cells into wild-type or PINK1-deficient mice. We provide evidence for the entry and persistence of such cells in the brain and show that this leads to levodopa-reversible motor dysfunctions and partial degeneration of the nigrostriatal DA system in both genotypes. These findings establish that brain entry of autoreactive CD8^+^ T cells is sufficient to drive nigrostriatal degeneration and parkinsonian motor deficits, providing the most direct support to date for the hypothesis that an adaptive immune attack plays a key role in PD-like neurodegeneration.

## Introduction

The cardinal motor symptoms of Parkinson’s disease (PD) are driven in part by the selective and progressive degeneration of dopamine (DA) neurons in the substantia nigra pars compacta (SNc), leading to striatal DA depletion and disruption of basal ganglia circuits controlling movement dynamics. Although DA replacement therapy can transiently restore motor function, no disease-modifying therapies currently exist. Genetic studies in familial forms of PD have implicated defects in fundamental cellular systems, such as in mitochondria, lysosomes and vesicular trafficking, arising from loss-of-function variants in genes such as *Pink1*, *Prkn* (Parkin), *Lrrk2* or *Gba* [1–9]. Such mutations lead to defects across multiple cell types, including neurons, but also immune cells [1, 10]. How these cellular defects translate into the selective neuronal vulnerability seen in PD represents a fundamental unanswered question.

Immune cells and inflammatory mechanisms are increasingly considered as playing a role in the initiation or progression of PD. Notably, cytokines such as IL-1β, TNF-α, IFN-γ, IL-6 and IL-8 are elevated in the serum, cerebrospinal fluid and nigrostriatal tissue from PD patients relative to age-matched controls [11–15]. The presence of T cells in the brain of PD subjects has also been reported [13, 16, 17] and T cells recognizing specific antigens like α -synuclein and PINK1 have been detected in the blood of PD patients [18–21]. Interestingly, overexpression of α-synuclein in mice leads to brain entry of CD4⁺ T-cells, neuroinflammation and DA neuron loss in mice and rats [22–25]. Immune mechanisms are also increasingly recognized as triggered by loss-of-function of PINK1 or Parkin, shown to disinhibit mitochondrial antigen presentation (MitAP) on MHC class I molecules in antigen-presenting cells (APCs) [1]. Strikingly, in *Pink1*^-/-^ mice, gastrointestinal infection leads to an increased abundance of mitochondrial antigen-specific CD8⁺ T cells in the periphery and the loss of striatal DA-terminal markers as well as L-DOPA-reversible motor deficits [26]. These findings support a paradigm shift suggesting that autoimmune mechanisms act as a prime driver of neuronal loss in some forms of PD [10, 27, 28].

A critical unanswered question is whether brain entry of autoreactive mitochondrial antigen-specific CD8⁺ T cells is sufficient to induce DA system dysfunction. To address this possibility, we took advantage of transgenic mice that express a T cell receptor (TCR) specific for a mitochondrial antigen derived from 2-oxoglutarate dehydrogenase (OGDH) [29, 30]. OGDH represents a uniquely relevant autoantigen in the context of PD. OGDH is a highly abundant, matrix-localized mitochondrial protein expressed in all cells including in DA neurons [31]. Importantly, OGDH-derived peptides are selectively presented on MHC class I molecules following mitochondrial stress or in absence of PINK1 or Parkin in antigen-presenting cells [1, 26]. This presentation could then induce the response of OGDH-specific CD8⁺ T cells which may then attack DA neurons upon entry in the central nervous system. We hypothesized that peripheral adoptive transfer of pre-activated mitochondrial antigen specific CD8⁺ T cells into *Pink1*^-/-^ mice would result in brain infiltration and targeted attack of the DA system. Compatible with this hypothesis, we report that such transfer leads to L-DOPA-reversible motor impairment and robust loss of SNc DA neurons and striatal axonal markers in both WT and *Pink1*^-/-^ mice. Such findings suggest that neuronal loss of PINK1 function in *Pink1*^-/-^ mice may not be a necessary contributor to the induction of neurodegeneration. This work provides a new perspective on immune mechanisms capable of driving PD pathology. Importantly, our observation of CD8^+^ T-cell-mediated loss of DA neurons also in WT mice raises the possibility that similar mechanisms could also be at play in sporadic forms of PD.

## Results

### Mitochondrial antigen-specific CD8^+^ T cells infiltrate the brain and are found in higher numbers in *Pink1*^-/-^ mice

We tested the hypothesis that mitochondrial antigen-specific CD8^+^ T-cells (thereafter called Mito-T) can induce PD-like pathology and parkinsonism in mice following their adoptive transfer into *Pink1*^-/-^ mice. To do so, we purified CD8^+^ T cells from 2C TCR transgenic mice, in which CD8^+^ cells express a TCR specific for the mitochondrial antigen OGDH and adoptively transferred 5 X 10^6^ *in vitro* activated CD8^+^ cells by intra-peritoneal (i.p.) injection into WT or *Pink1*^-/-^ mice (**Fig. 1A and Supplementary Fig. 1A**). As a control, WT and *Pink1*^-/-^ mice were adoptively transferred with activated CD8^+^ T cells (thereafter called OVA-T) from OT-I TCR transgenic mice, expressing a TCR specific for the ovalbumin antigen, not found in mice. Forty-eight hours after adoptive transfer, pertussis toxin (PTx) (20 µg/kg) was administered i.p. to facilitate brain infiltration of the transferred CD8^+^ T cells, as typically done in rodent multiple sclerosis models. Blood samples were collected on days 7 and 40 after adoptive transfer to assess the persistence and abundance of the transferred cells. The transferred cells were tracked using a monoclonal antibody (1B2) specific for the TCR expressed by the Mito-T CD8^+^ T cells or with the K^b^-OVA tetramer for control OVA-T CD8^+^ T cells. The gating strategy is described in **Supplementary Fig. 1B-C**. We successfully detected the transferred Mito-T and OVA-T cells in the blood of the recipient mice at both time points, with a higher frequency of the Mito-T compared to the OVA-T cells (**Fig. 1B-C)**. Although the proportion of Mito-T cells in the blood of the transferred mice was not significantly different between *Pink1*^-/-^ and WT mice (**Fig. 1B-C**), the absolute number of such cells detected in the spleen at day 40 was higher in the *Pink1*^-/-^ mice compared to WT littermates (**Fig. 1D**). Altogether, these observations indicate that Mito-T cells accumulate more than OVA-T cells in *Pink1*^-/-^ and WT mice, suggesting that such mitochondrial antigen specific CD8^+^ T cells encounter their antigen in recipient mice. Moreover, as more Mito-T CD8^+^ T cells were recovered in *Pink1*^-/-^ mice compared to WT recipients, these findings suggest that they may have seen more mitochondrial antigens in the peripheral tissues of *Pink1*^-/-^ mice.

**Figure 1:**
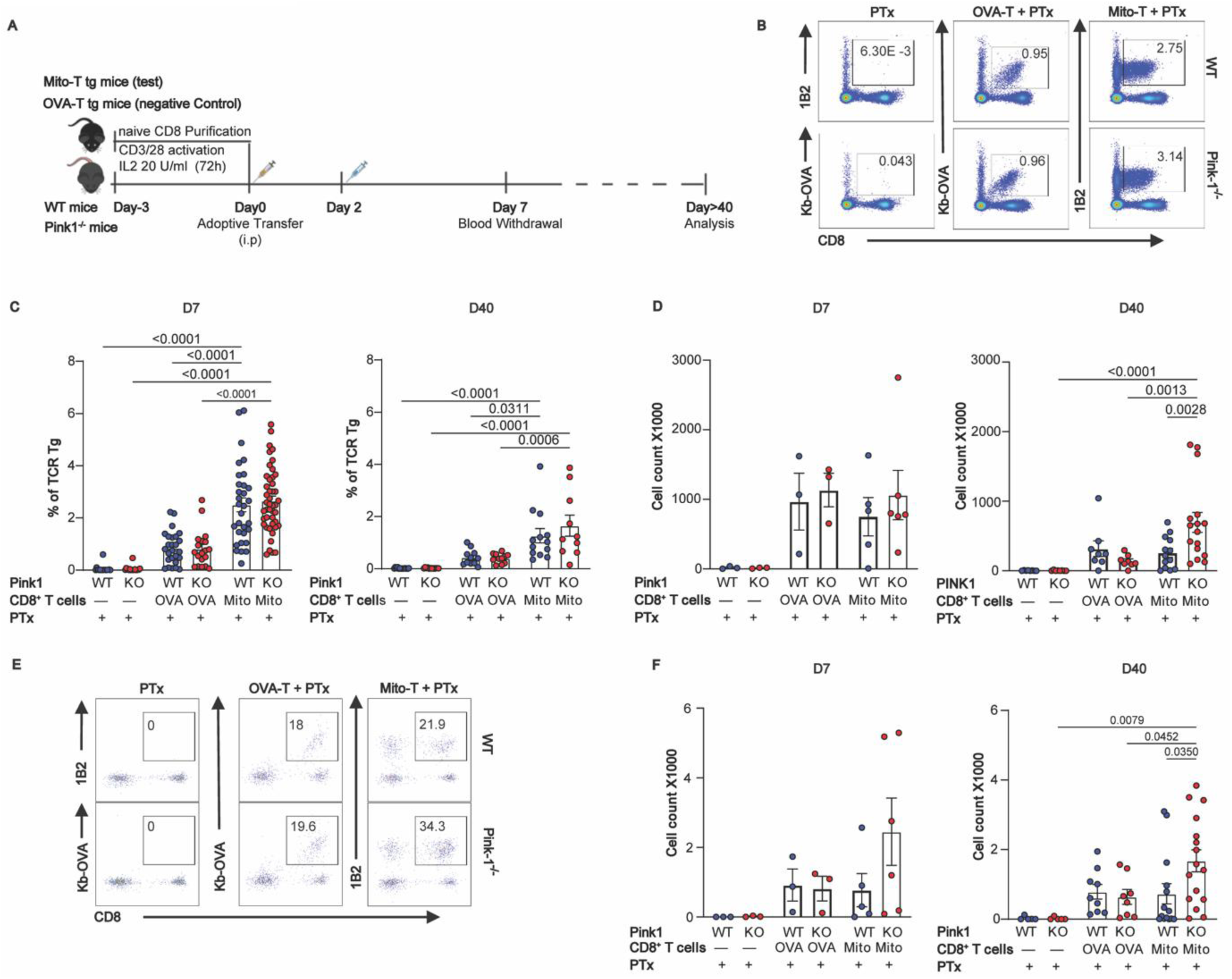
Adoptively transferred mitochondrial antigen-specific CD8^+^ T cells persist in the circulation and infiltrate the brain in higher amounts in Pink1^-/-^ mice. **A.** Schematic diagram outlining the experimental workflow. **B.** Representative flow cytometry profiles for the detection of adoptively transfer TCR transgenic CD8^+^ T cells. Mito-T cells were identified using the anti-TCR clonotype antibody 1B2 (CD8^+^1B2^+^) while OVA-T CD8^+^ T cells were identified using the K^b^-OVA tetramer (CD8^+^K^b^-OVA^+^). PTx refers to mice that were not adoptively transferred and only received the Ptx treatment. **C.** Frequency of the adoptively transferred TCR transgenic CD8^+^ T cells in the blood at day 7 and 40 after the adoptive transfer in WT and Pink1^-/-^ mice. D. Absolute number of adoptively transferred TCR transgenic CD8^+^ T cells in the spleen of WT and Pink1^-/-^mice. **E.** Representative flow cytometry profiles for the detection of adoptively transfer TCR transgenic CD8^+^ T cells that have infiltrated into the brain. **F.** Absolute number of adoptively transferred TCR transgenic CD8^+^ T cells that infiltrated the brain of WT and Pink1^-/-^ mice at 7 days (left) and 40 days (right) after the adoptive transfer. Each dot represents the results obtained from one mouse. Data is representative of a minimum of three independent experiments. p values were determined using two-way ANOVA followed by Tukey’s multiple comparisons post-hoc test. Data are presented as mean ± s.e.m. Data from 3 independent experiments, male and female mice were pooled.

We next asked whether the adoptively transferred Mito-T CD8^+^ cells were infiltrating the brain of the recipient mice. To assess brain infiltration, mice were administered FITC-conjugated anti-CD45 antibody intravenously 3–5 min before tissue collection to label circulating leukocytes but not parenchymal infiltrating cells (**Supplementary Fig. 1B-C**). Flow cytometry revealed the presence in the brain of both Mito-T and OVA-T cells at both time points, with higher abundance at day 40 (**Fig. 1E-F**) suggesting local antigen recognition and expansion of Mito-T cells. Strikingly, absolute counts of brain-infiltrating Mito-T cells were greater in *Pink1*^-/-^ mice than in WT littermates (**Fig. 1E-F**). Mito-T cell infiltration in the brain was also associated with an increase of recipient T cells (**Supplementary Fig. 1D**). The findings demonstrate that mitochondrial antigen-specific CD8^+^ T-cells infiltrate and persist within the brain for at least 40 days post-transfer, and that enhanced brain accumulation occurs in the absence of functional PINK1.

### Adoptive transfer of mitochondrial antigen-specific CD8^+^ T cells induces L-DOPA reversible motor impairment in both *Pink1*^-/-^ and WT mice

Our observation of entry of the adoptively transferred CD8^+^ T cells in the brain raised the possibility that Mito-T cells recognize their antigen in the brain, leading to DA neuron loss and motor circuit dysfunction. Compatible with this hypothesis, qualitative observations of the animals in their home cage revealed that after a period of 30-40 days following the adoptive transfer, some of the mice showed impaired locomotion. To validate this, we performed more extensive behavioral phenotyping using open field locomotor analysis, grip strength analysis, a rotarod task and the pole test. In the open field, we found reduced movement distance and vertical episodes in the mice transferred with Mito-T cells, but not in the mice transferred with OVA-T cells or only injected with PTx (**Fig. 2A-B**). Similar effects were seen in *Pink1*^-/-^ and WT recipients. Movement time (total ambulation + stereotypy) and average movement velocity were also reduced in the Mito-T condition, while the number of rest episodes did not differ between the groups (**Supplementary Fig. 2**). Time spent in the center of the open field (thigmotaxis), often used as an indirect index of anxiety, was not altered (**Fig. 2C**). Grip strength was also unaffected (**Fig. 2D**), suggesting preserved muscle force. In the rotarod task, the maximal speed leading to falling off the apparatus and the time to fall was lower in the mice that received Mito-T cells compared to the other groups (**Fig. 2E**), independently of genotype. Finally, in the pole test, the mice with adoptive transfer of Mito-T cells, but not the mice transferred with OVA-T cells or treated only with PTx showed impaired performance, as revealed by a slower time to descend (**Fig. 2F**, left panel), a deficit abolished by pre-treatment with the DA synthesis precursor L-DOPA (**Fig. 2F**, right panel), arguing that the motor impairments observed in the adoptively transferred mice were linked to reduced DA levels. Like for the other behavioral tests, *Pink1*^-/-^ and WT littermates did not differ, suggesting that a similar neurodegenerative process occurred in both genotypes.

**Figure 2:**
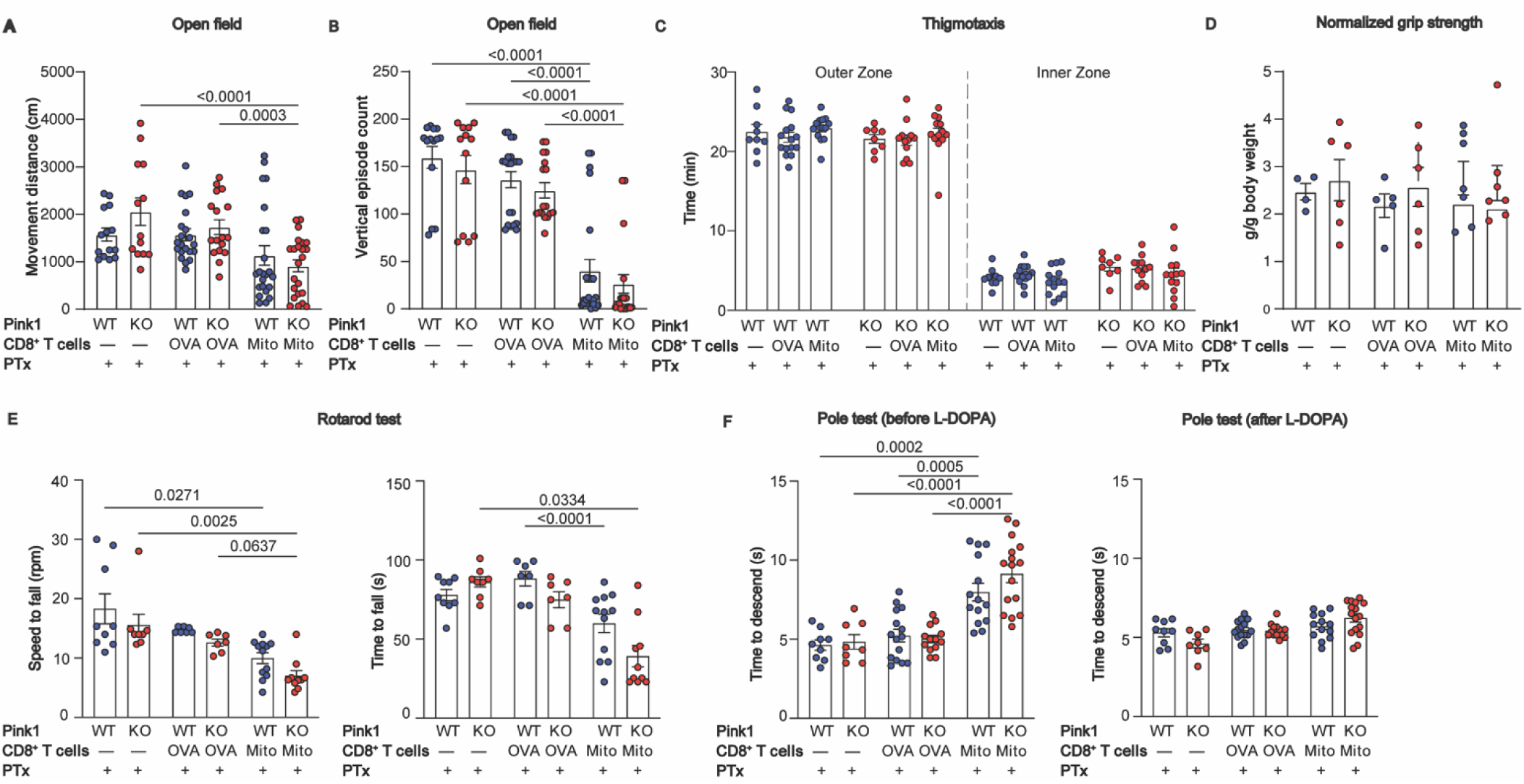
Adoptive transfer of mitochondrial antigen-specific CD8^+^ T cells leads to L-DOPA reversible motor impairment. Motor functions were analyzed between days 42 and 56 after the adoptive T cell transfer. **A.** In the open field test, the mice were examined for a period of 30min. The total movement distance travelled by the mice in an open field arena was measured. **B.** Vertical episodes (number of time that the mouse rears). **C.** Time spent in the outer zone vs inner zone of the open field. **D.** Grip strength for the four limbs normalized by mouse body weight. **E.** Rotarod analysis of the maximal speed and time to fall. **F.** Time to descend in the pole test before (left panel) and 15-30 min after i.p. administration of L-DOPA (25mg/kg) and the dopadecarboxylase inhibitor benserazide (6.5 mg/kg) (right panel). Data are representative of four independent experiments for A, B, C, and F and three independent experiments for D and E. p values were determined via two-way ANOVA followed by Tukey’s multiple comparisons test. Data are presented as mean ± s.e.m. The data combines both male and female mice.

### Adoptive transfer of mitochondrial antigen–specific CD8⁺ T cells induces degeneration of nigrostriatal dopamine neurons

The motor dysfunctions observed after adoptive transfer of Mito-T CD8⁺ T cells suggested impairment of basal ganglia circuitry, a system critically dependent on DA and known to be particularly vulnerable to cellular stress. Since DA neurons can express MHC class I and potentially present antigens to CD8⁺ T cells [32], we hypothesized that infiltration of Mito-T cells into the brain would directly compromise the nigrostriatal DA system. To test this, we performed immunohistochemical analyses of the striatum from mice 30-40 days post-transfer. In the dorsal striatum, the primary projection target of SNc neurons and the region most severely affected in PD, recipients of Mito-T cells exhibited marked reductions in tyrosine hydroxylase (TH) immunoreactivity. This decrease reached statistical significance in *Pink1*^⁻/⁻^ but not in WT mice, which showed more variable effects (**Fig. 3A-B**). No loss was observed in OVA-T cells recipients or in PTx-only controls. Importantly, the density of serotonin neuron axon terminals (SERT immunoreactivity) and the total number of intrinsic striatal neurons (NeuN) remained unchanged across all groups (**Figs. 3A-B**), suggesting that the degenerative process was selective for DA neuron projections. These findings suggest a direct CD8^+^ T cell attack of DA neurons, impacting the axonal domain, providing an explanation for the impaired behavioral performance of mice that received Mito-T CD8^+^ T cells.

**Figure 3:**
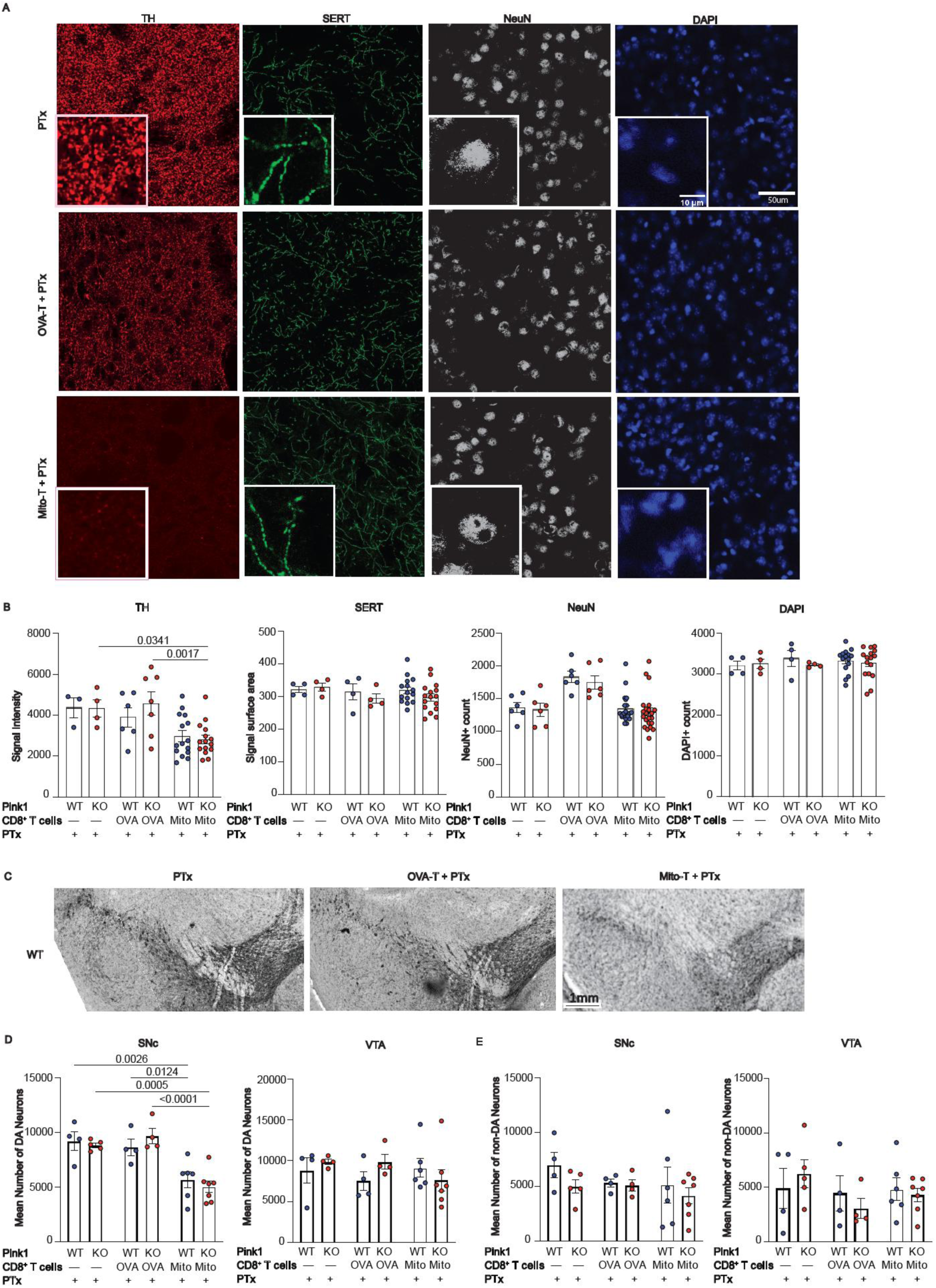
Adoptive transfer of mitochondrial antigen–specific CD8^+^ T cells cause selective dopaminergic denervation. **A.** Immunofluorescence staining of the dorsal striatum of Pink1^-/-^ and WT mice shows dopaminergic neuronal markers tyrosine hydroxylase (TH, red). Serotonergic terminals were visualized using the serotonin transporter (SERT, green), neuronal nuclei with NeuN (grey), and total nuclei with DAPI (blue). **B.** Quantification of axon terminal marker signal intensities revealed a selective reduction of TH but not SERT, NeuN or DAPI, consistent with selective dopaminergic denervation in Pink1^-/-^ mice after adoptive transfer of mitochondrial antigen–specific CD8^+^ T cells. NeuN and DAPI are represented as absolute cell counts per unit area. **C.** Representative images of the DAB staining used for stereological counting of TH^+^ DA neurons in the ventral midbrain. **D,E.** Stereological counting of TH^+^ (DA neurons) and non-DA neurons in the ventral tegmental area (VTA) and substantia nigra pars compacta (SNc). Data pooled from three independent experiments. Brains were collected 42–56 days post-transfer. Statistical significance was determined by two-way ANOVA followed by Tukey’s multiple comparisons test. Data are shown as mean ± s.e.m.

We next asked whether striatal TH loss reflected selective axon terminal degeneration or was instead a consequence of cell body loss in the ventral midbrain. Unbiased stereological counts of TH⁺ neurons revealed a significant reduction of DA neuron cell bodies in the SNc in recipients of Mito-T cells compared to OVA-T controls (**Fig. 3C-D**). In contrast, the number of VTA DA neurons was unchanged (**Fig. 3D**). The magnitude of SNc cell body loss was comparable in both *Pink1*^⁻/⁻^ and WT mice (**Fig. 3D**), and no differences were observed in the number of non-DA neurons in either region (**Fig. 3E**).

Microglia and astrocytes are increasingly considered to be involved in mediating neuroinflammatory and neurodegenerative processes in PD. For example, the number of microglia labelled with Iba1 is increased in many mouse PD models [33, 34]. We therefore next sought to determine whether the persistent presence of T cells in the mice after adoptive transfer was accompanied by sustained activation of resident glial populations. We examined brain sections at a sub-chronic phase, 30 days after adoptive transfer of Mito-T or OVA-T cells. Quantification of Iba1⁺ microglial immunostaining [35] in dorsal striatum revealed no significant differences in mean fluorescence intensity between Mito-T, OVA-T or control mice (**Supplementary figure 3A**). To confirm these histological findings, we performed Western blot analysis of dissected striata and SNc tissue blocks. The protein levels of Iba1 and of the astrocytic marker GFAP were not significantly altered in Mito-T or OVA-T mice compared with controls (**Supplementary figure 3B**).While these observations do not exclude the presence of functional changes in these cells, these results indicate that at this point, adoptive transfer of activated mitochondrial antigen specific CD8⁺ T cells does not induce broad or sustained upregulation of microglial or astrocytic markers in the striatum or SNc. Collectively, these results demonstrate that adoptive transfer of mitochondrial antigen–specific CD8⁺ T cells can trigger selective degeneration of nigrostriatal DA neurons, resulting in both loss of striatal terminals and degeneration of SNc cell bodies. Moreover, once T cells infiltrate the CNS, they elicit comparable SNc neuronal soma loss in both WT and *Pink1*-deficient mice.

### Adoptively transferred CD8⁺ T cells acquire a memory-like phenotype

The persistence of the adoptively transferred CD8⁺ T cells within the CNS for up to 40 days prompted us to examine whether they have undergone differentiation into memory T cells, providing a potential explanation for the lag time between the initial entry of the adoptively transferred cells in the brain and the appearance of motor dysfunctions and neuronal loss. Flow cytometric profiling of brain-infiltrating CD8⁺ T cells at day 40 post-transfer revealed that both Mito-T and control OVA-T cells acquired a memory phenotype. These cells exhibited high expression of CD44, CD127, and Ly6C, together with low expression of the exhaustion marker PD-1, consistent with long-lived, functionally competent memory T cells rather than terminally exhausted cells (**Fig. 4A–C**). Both Mito-T and OVA-T cells isolated from the brain expressed CXCR3 and CXCR6 providing an explanation for CNS tropism (**Fig. 4A**). The different subsets of memory, effector, resident or central T cells were all detected in the brain **(Fig. 4C).** These findings suggest Mito-T cells persist as memory CD8^+^ T cells, and even establish residency, in a phenotype poised for antigen recognition and effector function within the CNS. Notably, although Mito-T and OVA-T cells displayed largely overlapping phenotypic profiles, the total number of recovered brain-infiltrating cells differed between the two populations, indicating that antigen specificity influenced their accumulation or persistence within the CNS rather than their differentiation state.

**Figure 4:**
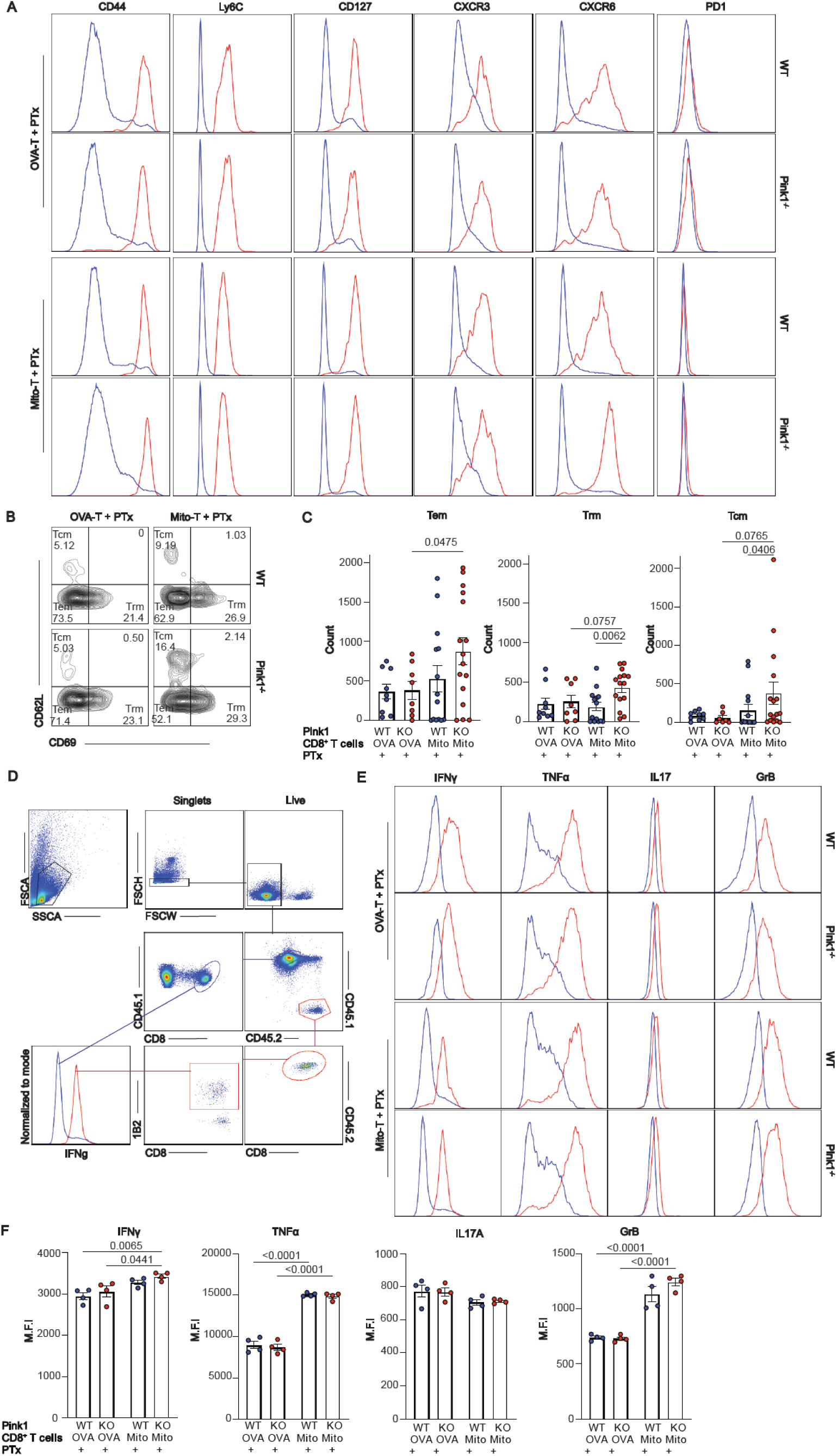
Brain-infiltrating CD8⁺ T cells acquire a memory phenotype and retain effector function following adoptive transfer. **A.** Representative flow cytometry plots showing expression of memory- and homing-associated markers (CD44, Ly6C, CXCR3, CXCR6, CD127, and PD-1) on brain-infiltrating CD8⁺ T cells isolated 40 days after adoptive transfer of OVA-T or Mito-T CD8⁺ T cells into WT or Pink1⁻/⁻ recipient mice treated with PTx (red) vs rest of CD45^+^cells in the brain (blue). **B.** Representative flow cytometry profile of CD69 and CD62L expression on brain-infiltrating CD8⁺ T cells, allowing discrimination of tissue-resident memory (Trm; CD69⁺CD62L⁻), effector memory (Tem; CD69⁻CD62L⁻), and central memory (Tcm; CD69⁻CD62L⁺) subsets. **C.** Quantification of the different memory CD8⁺ T cell subsets among OVA-T and Mito-T populations recovered from the brain of WT and Pink1⁻/⁻ mice. **D.** Flow cytometry gating strategy for assessment of effector function in brain-infiltrating CD8⁺ T cells. CD45.2⁺ donor-derived T cells isolated from the brain 30 days post-transfer were mixed with congenic CD45.1⁺ B6SJL splenocytes and stimulated ex vivo with PMA and ionomycin prior to intracellular cytokine staining. **E.** Representative flow cytometry plots showing intracellular expression of IFN-γ, TNF-α, GrB, and IL-17A in OVA-T and Mito-T CD8⁺ T cells (red) vs B6SJL CD8^+^ cells (blue) following stimulation. **F.** Quantification of mean fluorescence intensity (MFI) for IFN-γ, TNF-α, granzyme B, and IL-17A in donor-derived TCR⁺ CD8⁺ T cells. Data is representative of two independent experiments. p values were determined using two-way ANOVA followed by Tukey’s multiple comparisons test. Data are presented as mean ± s.e.m. Data from male and female mice were pooled.

A key complementary question is whether these brain-infiltrating CD8^+^ T cells are functional, i.e. producing cytokines and cytotoxic molecules. To tackle this question, Mito-T and OVA-T cells isolated from the brain 30 days after adoptive transfer were examined for the production of effector molecules (IFN-γ, TNF-α, IL-17 and granzyme B). As very few cells were recovered, the sorted cells (CD45.2^+^) were mixed with congenic B6SJL splenocytes (CD45.1^+^) (**Fig. 4D**) before *in vitro* stimulation with PMA and ionomycin for 5h to assess their effector functions. The brain-infiltrating memory CD8^+^ T cells produced high levels of IFN-γ, granzyme B, and TNF-α, whereas IL-17A production was almost non-detectable (**Fig. 4E-F**), consistent with a classical type 1 cytotoxic (Tc1) effector profile. The production of effector molecules was also observed in OVA-T cells as expected for cells with a memory phenotype. Altogether, our results demonstrate that the adoptive transfer of mitochondrial antigen specific CD8^+^ T cells leads to their accumulation as functional memory T cells in the brain. Their robust cytotoxic effector function suggests that they are endowed with the capacity to provoke neuronal injury, providing an explanation for the degeneration of DA neurons observed in the mice after adoptive transfer of Mito-T CD8^+^ T cells.

## Discussion

The origin of neuronal degeneration in PD is an unsolved mystery. A large body of work has examined and documented the possibility that the loss of DA neurons results from cell-autonomous dysfunctions in these neurons. This includes the hypothesis that pathological aggregation of the protein α-synuclein leads to cellular dysfunctions in DA neurons and the activation of apoptosis [36–38]. Or that aggregation of α-synuclein leads to reduced levels of monomeric forms of this protein, with loss of function mechanisms leading to neuronal demise [39, 40]. It has also been hypothesized that environmental exposure to some pesticides leads to the loss of DA neurons by blocking the mitochondrial complex 1 in these cells [41]. Partial support for this hypothesis comes from the observation that conditional deletion of mitochondrial complex 1 gene products in DA neurons leads to age-dependent gradual impairment of the DA system in mice [42–44].

An equally plausible but less examined alternate or complementary hypothesis is that the loss of SNc DA neurons results from dysfunctions in other cell types including glial cells in the brain or peripheral immune cells that gain access to the brain. The work described in the present manuscript adds strong support for such non-cell autonomous mechanisms by showing that brain entry of autoreactive mitochondrial-targeting CD8^+^ T cells is sufficient to cause the loss of DA neurons and to induce motor dysfunctions, a phenotype suggestive of PD pathology. In accordance with this hypothesis, a growing series of studies focusing on gene products linked to familial, genetic forms of PD, suggest that multiple PD-related proteins including Parkin, PINK1 and LRRK2 act as regulators of innate and adaptive immune responses in antigen presenting cells (APCs) and that loss of function of some of these proteins can amplify immune responses [1, 10, 18–20, 27, 28, 45–47]. Recent work revealed that in inflammatory conditions, PINK1 and Parkin restrict the presentation on MHC class I molecules of antigens derived from proteins present in the mitochondrial matrix (MitAP) [1]. This led to the hypothesis that a loss of function of PINK1 or Parkin leads to increased presentation of mitochondrial antigens and to the activation of autoreactive CD8^+^ T cells [1]. In keeping with this hypothesis, follow-up work revealed that MitAP can be activated following gut infection with Gram-negative bacteria in *Pink1*^-/-^ mice, a process associated with the activation and expansion of mitochondrial antigen-specific CD8^+^ T cells and the concomitant development of motor impairments reversible by L-DOPA [26]. Although thought-provoking, a direct role for CD8^+^ T cells in causing neuronal loss and the emergence of motor impairments was not clearly established in this previous work.

The present study provides direct evidence that mitochondrial antigen-specific CD8⁺ T cells alone are sufficient to trigger DA neuron loss and L-DOPA-reversible motor symptoms. Beyond supporting the hypothesis that MitAP and CD8^+^ T cells are possible initiators of PD-like pathology, our work delineates a clear periphery-to-CNS immune axis where primed autoreactive CD8⁺ T cells enter the brain, establish a tissue-resident memory phenotype and induce neurodegeneration. This positions mitochondrial autoimmunity as a mechanistic bridge between systemic immune dysregulation and selective neuron loss.

Although the adoptive transfer of pre-activated antigen-specific CD8⁺ T cells represents a reductionist experimental approach, this strategy was deliberately designed to isolate and test a single unresolved causal question: whether autoreactive mitochondrial-specific CD8⁺ T cells are sufficient to induce selective DA neuron degeneration once they gain access to the brain. By removing confounding variables such as infection, α-synuclein overexpression, or generalized inflammation, this model reveals the intrinsic neurodegenerative potential of mitochondrial antigen-specific CD8⁺ T cells.

A surprising finding in the present work is that neurodegeneration and motor impairments developed in both *Pink1*^-/-^ and WT mice following mitochondrial-targeted T cell adoptive transfer. Although further work will be required to clarify this, a possibility is that since PINK1 inhibits the presentation of mitochondrial antigens by APCs, one of the main roles of this protein in the context of PD is to prevent the initiation of the disease process by restricting the priming of naive CD8^+^ T cells in the periphery [48]. Accordingly, introducing preactivated cytotoxic T cells by adoptive transfer bypasses the protective function of PINK1 and directly engages a pathophysiological process in both WT and KO mice, as long as some level of MitAP occurs in DA neurons or local glial cells, allowing the autoreactive CD8^+^ T cells to engage with brain cells. This scenario implies that loss of function of PINK1 in DA neurons may perhaps not be required to engage neuropathological mechanisms in PD once cytotoxic T cells are generated in the periphery. Our work thus provides a different perspective on how PINK1 loss of function could lead to dysfunction and loss of DA neurons. PINK1 was first shown to be involved in the clearance of damaged mitochondria by mitophagy [49–53]. Such observations were taken by some to imply that loss of PINK1 function in DA neurons would lead to the accumulation of damaged mitochondria in these neurons, eventually promoting their death. However, mitophagy still proceeds in the absence of PINK1 [54–57] and there is presently very limited evidence that dysfunctional mitochondria indeed accumulate in PINK1 or Parkin KO DA neurons [58]. Although we did not directly evaluate the functions of neuronal PINK1 in the present study, our results are more compatible with the hypothesis that loss of function of PINK1 engages autoimmune mechanisms in the periphery at early stages of the disease process. Our finding of loss of DA neurons even in WT mice raises the thought-provoking hypothesis that the T-cell-mediated neuronal death mechanism identified in the present model could occur not only in individuals carrying mutations in PINK1, but also in sporadic forms of PD. One can hypothesize that age-related decreases in the levels of PINK1 or Parkin [59] in APCs could engage a sequence of events similar to that induced by loss of function mutations.

The specific cellular and molecular mechanisms by which CD8^+^ T cells kill DA neurons in the present model remains to be determined. Based on previous work suggesting that DA neurons have the capacity to express MHC class I molecules on their surface [32], one possibility is that in these mice, following entry in the brain of Mito-T cells, DA neurons present mitochondrial antigens leading to their direct recognition and attack by cytotoxic CD8^+^ T cells. However, more work is needed to characterize the extent and dynamics of MHC class I expression by DA neurons in both *Pink1*^-/-^ and WT mice. Our finding of an equivalent attack of DA neurons by activated Mito-T CD8^+^ T cells in the two genotypes suggests that a minimally sufficient level of antigen presentation may happen in both genotypes, although not necessarily at the same levels. Further work will be required to clarify this and to examine the alternate possibility that MitAP occurs principally in glial cells, leading to the production of neurotoxic signals by these cells or to reduced production of neurotrophic signals. A growing literature suggests similar mechanisms implicating CD8^+^ T cells and glial cells in Alzheimer’s disease [49, 60–62].

Our findings further demonstrate that brain-infiltrating antigen-specific CD8⁺ T cells retain robust effector capacity, as evidenced by their ability to produce IFN-γ, TNF-α, and granzyme B upon *ex vivo* stimulation. This effector profile indicates that infiltrating CD8⁺ T cells are not functionally exhausted or inert but instead remain poised to mediate inflammatory and cytotoxic responses within the CNS. The presence of granzyme B together with pro-inflammatory cytokines supports a model in which CD8⁺ T cells may contribute to neuronal injury through both direct cytotoxic mechanisms and indirect amplification of neuroinflammation.

Taken together, our observations provide unprecedented support for a key role of mitochondrial antigen-specific CD8^+^ T cells in DA system perturbations and associated parkinsonism in a mouse model of early-onset PD. This work could be instrumental for the development of better animal models of PD and for the identification of new immune-based therapeutic approaches to treat PD patients.

## Acknowledgements

We would like to thank Drs. Michel Desjardins, Samantha Gruenheid, Heidi McBride and Janelle Drouin-Ouellet for their critical feedback during the development of this project. We also thank Dr. Numa Dancause for providing access to their stereological counting microscope system. We thank Frédéric Duval and Marie-Josée Aubin for cell sorting and the animal care technicians for mice husbandry. The present study was funded by the joint efforts of the Michael J. Fox Foundation for Parkinson’s Research (MJFF) and the Aligning Science Across Parkinson’s (ASAP) initiative. MJFF administers the grant ASAP 000525 on behalf of ASAP and itself. The Trudeau lab also received support from the Canadian Institutes of Health Research (CIHR) (grant 165928) and from the Krembil foundation. M. N. E. received a graduate student award from the mission sector of the ministry of higher education, Egypt.

## Materials and methods

### Mice

The 2C TCR-transgenic mice (RRID:MGI:3621454) express a TCR recognizing an OGDH mitochondrial peptide presented by MHC class I K^b^ [29, 30], The OT-I TCR-transgenic mice (RRID:IMSR_JAX:003831) express a TCR specific for the OVA^257–264^ peptide presented by MHC class I K^b^ [63]. Male and female *Pink1*^-/-^ mice (Jackson Laboratory, Strain #017946, RRID: IMSR_JAX:017946) and WT littermate controls were bred and housed under specific pathogen-free conditions. All mice were maintained in accordance with the Canadian Council on Animal Care guidelines, under protocols approved by the animal ethics committee (Comité de déontologie de l’expérimentation animale, CDEA) of the Université de Montréal and of the Research Center of the Maisonneuve-Rosemont Hospital.

### T cell purification and activation

Naïve CD8^+^ T cells were isolated from the spleen of 2C and OT-I TCR transgenic mice using a Easysep^TM^ mouse naïve CD8^+^ T-cell isolation kit (STEMCELL, Technologies, cat# 19858). Purified naïve CD8^+^ T cells were then activated using plate-bound anti-CD3 antibody (clone 14-2C11, Bio X Cell, cat# BE0001-1, RRID: AB1107634) and a soluble anti-CD28 antibody (clone 37.51, Bio X Cell, cat# BE0015-1, RRID: AB1107624). Briefly, 24-well plates were coated with 1 µg/ml of anti-CD3 antibody in sterile PBS and incubated for 2h at 37°C, 5% CO2. 2 x 10^6^ naïve CD8^+^ T cells were seeded per well, and anti-CD28 antibody was added to a final concentration of 5 µg/ml. After 24h, recombinant mouse IL-2 (50 IU/mL, PeproTech, cat# 212-12) was added, and cells were maintained for 72h at 37 °C and 5 % CO₂. Activation was confirmed by flow cytometry analysis of up-regulation of CD44, CD25 and CD69 and down-regulation of CD62L.

### Adoptive transfer of TCR transgenic T cells

*Pink1*^-/-^ mice and their WT littermate controls were injected intraperitoneally (i.p) with 5 X 10^6^ *in vitro*-activated mitochondrial OGDH peptide specific CD8 T cells (Mito-T) derived from 2C TCR mice or OVA-257-264 peptide specific CD8^+^ T cells (OVA-T) derived from OT-I TCR mice, as a negative control. Cells were resuspended in 200 μl of sterile PBS immediately prior to injection. Control groups included mice receiving PBS but no T-cells. Forty-eight hours after adoptive transfer, all experimental animals received pertussis toxin (PTx; 20 µg/kg i.p., Cayman Chemical, cat# 23221) to facilitate T-cell entry into the central nervous system by transiently increasing blood–brain barrier permeability.

### FACS analysis

Blood samples (200 μl) were collected from the jugular vein into 100ul PBS containing 2mM EDTA at both early (day 7) and late (day 40) time points following the adoptive transfer. Spleens were harvested at day 7 from a subset of mice and from all the mice at the experimental endpoint. Red blood cells were lysed in 0.86% ammonium chloride (NH_4_CL) solution for 5 min at room temperature, followed by washing in PBS. Cell viability was assessed with Zombie NIR™ fixable viability dye (1:1000 in PBS; BioLegend, cat# 423105, RRID: AB_2565394) for 20 min at room temperature in the dark. Samples were then blocked with anti-CD16/CD32 (Fc Block; clone 2.4G2, BD Biosciences, cat# 553142, RRID: AB_394657) for 20 min at room temperature prior to surface staining. Surface marker staining was performed for 20 min at 4 °C in FACS buffer (Dulbecco’s PBS without phenol red supplemented with 3 % heat-inactivated fetal bovine serum, 10 mM HEPES, and 0.1 % sodium azide).

To identify OVA-T cells, samples were stained with H-2K^b^- OVA-257-264 peptide tetramer (NIH Tetramer core facility) and staining was performed at 37°C for 15 min at 37 °C before the antibody cocktail. Data acquisition was performed on a BD LSRFortessa™ X-20 flow cytometer (BD Biosciences, RRID:SCR_023638), and analysis was performed using FlowJo™ (BD Biosciences, RRID:SCR_008520). All antibodies (with clone, fluorochrome, manufacturer, catalog number, and RRID) are listed in **Supplementary Table 1**.

### Intracellular staining and effector function analysis

To assess effector function of brain-infiltrating CD8⁺ T cells, intracellular cytokine and granzyme B stainings were performed following short-term *ex vivo* stimulation. Brain-infiltrating leukocytes were isolated as described below followed by cell sorting of CD8^+^CD45.2^+^ cells. Due to the low yield of infiltrating T cells, sorted CD8^+^CD45.2⁺ T cells were mixed with congenic CD45.1⁺ B6.SJL splenocytes as carrier cells prior to stimulation. Cell suspensions were stimulated for 5h at 37 °C and 5 % CO₂ with phorbol 12-myristate 13-acetate (PMA, 50 ng/ml; Sigma-Aldrich) and ionomycin (1 µg/ml; Sigma-Aldrich) in complete RPMI-1640 medium. Brefeldin A (5 µg/ml; BioLegend) and monensin (2 µM; BioLegend) were added for the duration of the stimulation to block cytokine secretion. Following stimulation, cells were washed and stained with Zombie NIR™ fixable viability dye, followed by Fc receptor blocking (anti-CD16/32) and surface staining for CD45.1, CD45.2, CD8α, CD44 and TCR (1B2 or K^b^-OVA). Cells were then fixed and permeabilized using the Cyto-Fix/Cyto-Perm™ kit (BD Biosciences) according to the manufacturer’s instructions. Intracellular staining was performed for interferon-γ (IFN-γ), tumor necrosis factor-α (TNF-α), interleukin-17A (IL-17A), and granzyme B. Samples were acquired on a BD LSRFortessa™ X-20 flow cytometer, and data were analyzed using FlowJo™ software. Effector molecule production was quantified within viable CD45.2⁺CD8⁺1B2^+^ (for Mito-T) or CD45.2⁺CD8⁺K^b^-OVA^+^ (for OVA-T) T cells after exclusion of CD45.1⁺ carrier splenocytes. Unstimulated controls were included to define background staining and gating thresholds.

### Isolation of brain infiltrating lymphocytes

To distinguish infiltrated brain T cells from those still in circulation, mice were injected intravenously (i.v) with anti-CD45-FITC antibody (clone 30-F11, BD Biosciences, cat# 553080, RRID: AB_394609) 3–5 min before sacrifice for brain harvest. T cells shielded from this brief intravascular staining were considered to be localized within the brain parenchyma and were detected as CD45-FITC negative during flow cytometry analysis. Brains were dissected and minced longitudinally into ≥8 fragments using a sterile scalpel, then enzymatically digested in RPMI supplemented with 0.01 mg/ml DNase I (Roche, cat# 11284932001, RRID:SCR_019337) and 0.5 mg/ml collagenase D (Roche, cat# 11088858001, RRID:AB_2827424) for 40 min at 37 °C using the gentleMACS™ Octo Dissociator (Miltenyi Biotec, cat# 130-096-427). Cell suspensions were passed through a 70 µm cell strainer (Corning, cat# 352350) to remove debris. For lymphocyte enrichment, the digested tissue was centrifuged at 1500 rpm for 10 min, resuspended in 3 ml of 37% Percoll (Cytiva, cat# 17-0891-01) diluted in RPMI-1640, overlaid onto 3 ml of 70% Percoll, and centrifuged at 2000 rpm for 20 min at 25 °C without brake. The brain-infiltrating leukocyte fraction was collected from the interphase for downstream flow cytometry analysis.

### Immunohistochemistry

*Pink1*^⁻/⁻^ and WT littermate mice that had received adoptive transfer of CD8⁺ T-cells or control treatments were deeply anaesthetized with pentobarbital (70 µg/g body weight, i.p.; Wisent Bioproducts, cat# 450-166-XL) and perfused intracardially with 50 ml ice-cold PBS (0.01 M) followed by 50 ml ice-cold 4 % paraformaldehyde (PFA).

Brains were collected, post-fixed for 24h at 4°C in 4% PFA, then cryoprotected for 48h in 30% sucrose prepared in phosphate buffer (5.382g/l sodium phosphate monobasic, 8.662g/l sodium phosphate dibasic anhydrous; Sigma-Aldrich). Serial coronal free-floating sections (40μm) were obtained using a Leica CM1950 cryostat and stored in antifreeze solution (1.57g/l sodium phosphate monobasic anhydrous, 5.45g/L sodium phosphate dibasic anhydrous, 300ml ethylene glycol, 300ml glycerol, distilled water to volume) at −20 °C. Sections were permeabilized in 0.1% Triton™ X-100 (Sigma-Aldrich, cat# T8787) and blocked in 10 % BSA (Sigma-Aldrich, cat# A2153), containing 0.02 % sodium azide, 0.3 % Triton X-100, and 5% goat serum for 1h at room temperature. Overnight incubation (4 °C) was performed with primary antibodies: rabbit anti-TH (1:2000, AB152; Millipore, cat# AB152, RRID:AB_390204), rabbit anti-Sert (1:2000; Millipore, cat# PC177L-100UL, RRID:AB_10697452) or mouse anti-NeuN (1:1000; Abcam, cat# ab134014, RRID:AB_2313589). Nuclei were counterstained with DAPI (1:2000; Sigma-Aldrich, cat# D9542).

For the quantification of TH immunoreactivity in the striatum, three brain sections from each mouse were acquired from bregma 1.1, 0.14 and -0.94 and images from regions of interest including the whole striatum were obtained from both hemispheres and acquired using a 20X, 0.8 NA, plan apochromat objective on a Zeiss Axioscan 7 slide scanner (Zeiss Canada). Scans from whole striata were acquired for TH under identical exposure and illumination settings across all experimental groups to ensure quantitative comparability. TH terminal density was estimated by quantifying mean signal intensity across analyzed striatal fields. In separate experiments aimed at quantifying SERT, NeuN, DAPI or Iba1, images were acquired on a Nikon AX-R automated point-scanning confocal microscope (Nikon Instruments, RRID:SCR_023799) at 2048×2048 px resolution, Nyquist sampling, galvanometer scanning mode, with 1x averaging and 0.8ms dwell time, using a Plan Apo λ 40×/0.95 OFN25 DIC N2 objective lens (0.6 NA, Nikon Canada). For these quantifications, images were acquired from dorsal striatum fields, defined as the area superior to the anterior commissure and inferior to the corpus callosum. For each striata, 2-4 random fields per hemisphere were imaged. Quantification was performed in FIJI/ImageJ (NIH, RRID:SCR_002285) by applying a uniform background-subtracted threshold to all analyzed images. SERT+ terminal density was calculated as mean signal area across eight fields per dorsal striatum, with 5 sections/mouse. The number of NeuN-positive and DAPI-positive cells was quantified from the same image sets.

### Stereological analysis

Coronal brain sections were rinsed in 0.01 M PBS (1×) for 10 min and immersed in 0.01 M PBS containing 0.9 % H₂O₂ for 10 min to quench endogenous peroxidase activity. Sections were rinsed 3× in PBS (10 min each) and incubated for 48h at 4 °C with rabbit anti TH antibody (1:1000; Millipore, cat# AB152, RRID:AB_390204) diluted in 0.01 M PBS containing 0.3 % Triton X 100 (Sigma Aldrich, cat# T8787) and 50 mg/ml bovine serum albumin (BSA; Sigma Aldrich, cat# A2153). Coronal brain sections were rinsed in 0.01 M PBS (1X) for 10 min before immersed in 0.01 M PBS containing 0.9% H_2_O_2_ for 10 min to quench endogenous peroxidase activity. Sections were rinsed three times in 0.01 M PBS (10 min each) after which they were incubated for 48h at 4 °C with rabbit anti-TH antibody (1:1000; Millipore, cat# AB152, RRID:AB_390204) diluted in 0.01 M PBS containing 0.3 % Triton X 100 (Sigma Aldrich, cat# T8787) and 50 mg/ml bovine serum albumin (BSA; Sigma Aldrich, cat# A2153). Sections were rinsed 3× in PBS and incubated for 12h at 4 °C with biotin conjugated AffiniPure IgG (1:200; Jackson ImmunoResearch, cat# RON1231, RRID: AB_2337238). After washing 3× in PBS, sections were incubated for 3h at room temperature in streptavidin HRP conjugate (1:200; Cytiva/GE Healthcare, cat# RPN1231) diluted in 0.3 % Triton X 100/PBS. Visualization was performed using a 3,3′-diaminobenzidine tetrahydrochloride DAB)–glucose oxidase reaction (Sigma-Aldrich, cat# D8001) for 5 min. Sections were mounted onto charged microscope slides (Thermo Scientific, cat# J1800AMNZ) in 0.1 M acetate buffer, counterstained with cresyl violet (Sigma Aldrich, cat# HT901), dehydrated in graded ethanol, cleared in xylene, and finally cover slipped with Permount™ (Fisher Scientific. Cat # SP15-100). Unbiased stereological counts of TH⁺ neurons in the SNc and VTA were performed using the optical fractionator method [64] using Stereo Investigator v6 (MBF Bioscience, RRID:SCR_002526) at 100× magnification, with a 60 × 60 µm² counting frame, a 10 µm optical dissector, and two 1 µm guard zones. Sections were sampled every sixth slice, with counting sites placed at 100 µm intervals after a random start. A minimum of 6 sections per mouse containing the SNc was set as an inclusion criterion.

### Behavioral tests

All behavioral experiments were conducted during the dark (active) phase of the light/dark cycle, with mice maintained on a 12h light/12 h dark schedule (lights off 22:00–10:00). The experimenter was blinded to both genotype and the treatment group throughout all testing procedures. To minimize handling stress, mice were habituated to the experimenter for not less than 5 min per mouse on 3-5 consecutive days prior to testing. Mice were group housed (2-4 per cage) and tested 42-52 days post adoptive transfer. On testing days, mice were moved to the behavioral testing room and allowed to acclimate for 60 min before the start of assay.

### Basal locomotor activity

Basal locomotor activity was assessed using open field automated activity chambers (Superflex, Omnitech Electronics, Columbus, OH, USA) to quantify horizontal activity and vertical rearing episode. Mice were placed individually in clear acrylic activity arenas (41 × 41 cm) equipped with arrays of infrared photocell beams to detect movement. Horizontal and vertical activity was recorded continuously over 30 min by counting beam interruptions. Data were analyzed using Fusion software v5.3 (Omnitech Electronics, RRID:SCR_022276).

### Grip strength test

Muscular strength was measured with a grip strength meter (model BIO GS3, Bioseb, Vitrolles, France). Each mouse was weighed immediately prior to testing to allow normalization of strength data to body weight. For each trial, the mouse was gently allowed to grasp the metal grid with all four paws, and the peak force (g force) required to dislodge the animal was recorded. Three consecutive trials were performed per mouse, and the average peak force was calculated for analysis. Results are presented as gram force normalized to body weight (g/g).

### Pole test

The pole test apparatus consisted of a 60 cm high metal rod (10 mm diameter) wrapped in adhesive tape to improve grip, mounted securely in a test cage. For each trial, mice were placed head up at the top of the pole. Two parameters were recorded: (1) time to orient downward and (2) total time to descend until the forepaws reached the cage floor. Mice were first tested drug naïve and then re-tested 15–30 min after i.p. administration of either benserazide hydrochloride (6.5 mg/kg; Sigma Aldrich, cat# B7283) combined with L DOPA (25 mg/kg; Sigma Aldrich, cat# D1507) or an equal volume of sterile saline. A maximum cut off time of 180 s was imposed; mice exceeding this limit were scored as failures and returned to their home cage. Such mice were re-tested after a rest period.

### Western Blot Assay

Total protein lysates from brain tissue were prepared using Pierce™ RIPA Lysis Buffer (Thermo Scientific, cat# 89900) supplemented with complete™ EDTA free protease inhibitor (1 tablet/50 mL; Roche, cat# 11873580001) and PhosSTOP™ phosphatase inhibitor (Roche, cat# 04906837001). Protein concentrations were determined with the Pierce™ BCA Protein Assay Kit (Thermo Scientific, cat# 23227), and 20 µg total protein per sample was loaded for SDS PAGE. Proteins were separated on Mini PROTEAN® TGX™ precast gels (Bio Rad, cat# 456 1094) at 200 V for 40 min and transferred to Immobilon PVDF membranes (MilliporeSigma, cat# IPVH00010) at 100 V for 90 min. Membranes were blocked in PBST (PBS + 0.4 % Tween 20) containing 5 % non-fat dry milk for 30 min at room temperature, then incubated overnight at 4 °C with primary antibodies diluted in PBS containing 1 % bovine serum albumin (BSA; Sigma Aldrich, cat# A2153). Following 3 × 15 min washes in PBST, membranes were incubated with the appropriate HRP conjugated secondary antibodies (Jackson ImmunoResearch, various) in PBST + 5 % milk for not less than 30 min at room temperature, washed again 3 × 15min in PBST, and developed using Clarity™ Western ECL Substrate (Bio Rad, cat# 170 5061). Chemiluminescence was visualized with a ChemiDoc™ MP Imaging System (Bio Rad, RRID:SCR_019037).

### Statistical Analysis

Statistical analyses were performed using GraphPad Prism v10.0 (GraphPad Software, RRID:SCR_002798). The Mann–Whitney U test was used for 2 group comparisons; one way or two-way ANOVAs with Tukey’s post hoc test were used to compare 3 or more groups, as appropriate. Significance was set at p < 0.05. Data are presented as mean ± SEM unless otherwise specified.

## Data and materials availability

All key data is included in the manuscript. Access to original raw images is available on request from the corresponding authors.

