## Supplementary materials for "Mitochondrial antigen-specific CD8⁺ T cells drive dopamine neuron neurodegeneration"

[1]Department of neurosciences, Faculty of Medicine, Université de Montréal
[2]Neural Signaling and Circuitry research group (SNC)
[3]Center for Interdisciplinary Research on the Brain and Learning (CIRCA)
[4]Aligning Science Across Parkinson's (ASAP) Collaborative Research Network, Chevy Chase, MD, USA, 20815.
[5]Medical Biotechnology Department, Biotechnology Research Institute, National Research Centre, Dokki, Giza, Egypt
[6]Centre de recherche de l'hôpital Maisonneuve-Rosemont (CRHMR)
[7] Courtois Institute for Biomedical Innovation (Ci2B)
[8]Department of pharmacology and physiology, Faculty of Medicine, Université de Montréal
[9]Institut de recherches cliniques de Montréal (IRCM)
[10]Department of microbiology, infectious diseases and immunology, Faculty of Medicine, Université de Montréal
[11]Research Center of the Université de Montréal Hospital (CRCHUM)
[12]Dept. of Neuroscience and Neurology, Montreal Neurological Institute-Hospital, McGill University, Montreal,
[13]Department of medicine, Faculty of Medicine, Université de Montréal

### Author contributions

Conceptualization: N.L., D.M. and L-E.T.
Methodology: M.N.E., J-F.D., A.T., S.B., N.G., A.E.
Investigation: M.N.E., J-F.D., A.R., S.B., S.M., N.G., A.E, C.B.
Supervision: N.L., J.A.S. and L-E.T.
Writing—original draft: M.N.E., N.L. and L-E.T.
Writing—review & editing: M.N.E., N.L. and L-E.T.

**Running Title:** CD8 T cells drive dopamine neuron loss

**Corresponding authors:**

Dr. Louis-Éric Trudeau
Department of pharmacology and physiology
Faculty of Medicine
Université de Montréal

514-343-5692

Dr. Nathalie Labrecque
Institut de recherches cliniques de Montréal
Department of medicine, Faculty of medicine
Université de Montréal

514-987-550

**Extended Data:**

**Supplementary Table 1: List of antibodies used for flow cytometry.**

| **Antibody** | | **Supplier** | **Product information** | **RRID** |
|---|---|---|---|---|
| **1B2** | Biotin | Custom made | F23.1, 53-6.72 (rat anti-mouse CD8). | N/A |
| **CD11b** | BV711 | Biolegend | Clone M1/70; cat # 101242 | AB_2563310 |
| **CD127** | BV421 | Biolegend | Clone A7R34, cat # 135027 | AB_2563103 |
| **CD25** | APC | Biolegend | Clone PC61; cat # 102012 | AB_312861 |
| **CD4** | BV605 | Biolegend | Clone RM4-5; cat # 100548 | AB_2563054 |
| **CD44** | APC-cy7 | Biolegend | Clone IM7; cat # 103028 | AB_830785 |
| **CD45.2** | FITC | Biolegend | Clone 104; cat # 110706 | AB_313495 |
| **CD45.2** | Alexa flour 700 | Biolegend | Clone 104; cat # 109822 | AB_493731 |
| **CD62L** | PercP | Biolegend | Clone MEL; cat # 104430 | AB_2187124 |
| **CD69** | APC | Biolegend | Clone H1.2F3; cat # 104513 | AB_492844 |
| **CD8** | BV785 | Biolegend | Clone 53-6.7; cat # 100750 | AB_2562610 |
| **CXCR3** | PE | Biolegend | Clone CXCR3-173; cat # 126505 | AB_1027656 |
| **CXCR6** | PEdazzle594 | Biolegend | Clone SA051D1; cat # 151116 | AB_2721699 |
| **IFNγ** | PE | Biolegend | Clone XMG1.2; cat # 505807 | AB_315401 |
| **TNFα** | APC | Biolegend | Clone MP6-XT22; cat # 506307 | AB_315427 |
| **IL-17A** | APC | Biolegend | Clone TC11-18H10.1; cat # 506915 | AB_536016 |
| **Granzyme B** | PE | Biolegend | Clone QA16A02; cat # 372207 | AB_2563043 |
| **$K^b$-$OVA_{257-264}$** | | NIH Tetramer core facility | | N/A |

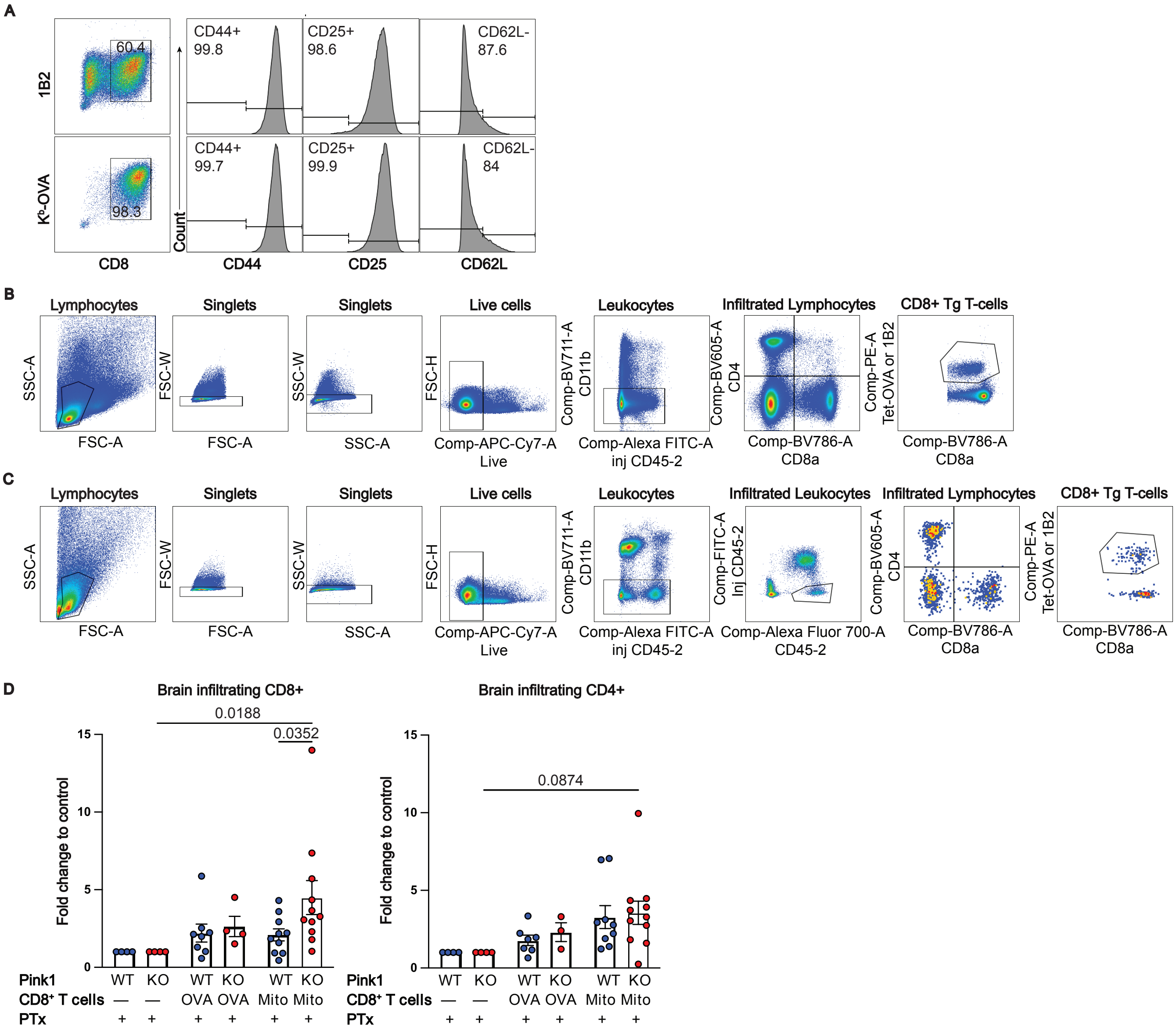


**Supplementary Figure 1. Adoptively transferred mitochondrial antigen-specific CD8$^+$ T cells persist in circulation and infiltrate the brain in higher amounts in *Pink1*$^{-/-}$ mice. A.** Representative flow cytometry chart for gating strategy used to detect the adoptively transfer TCR transgenic CD8$^+$ T cells in the spleen. **B.** Gating strategy to identify the brain infiltrating TCR transgenic CD8$^+$ T cells. **C.** Activation profile of OVA-T and Mito-T cells before the adoptive transfer. **D.** Fold change of endogeneous CD8$^+$ (right) and CD4$^+$ (left) brain infiltrating T lymphocytes, at day 40 post adoptive CD8$^+$ T cell transfer, compared to PTx genotype matched control. Data is representative of a minimum of three independent experiments. p values were determined using two-way ANOVA followed by Tukey's multiple comparisons test. Data are presented as mean ± s.e.m. Data from male and female mice were pooled.

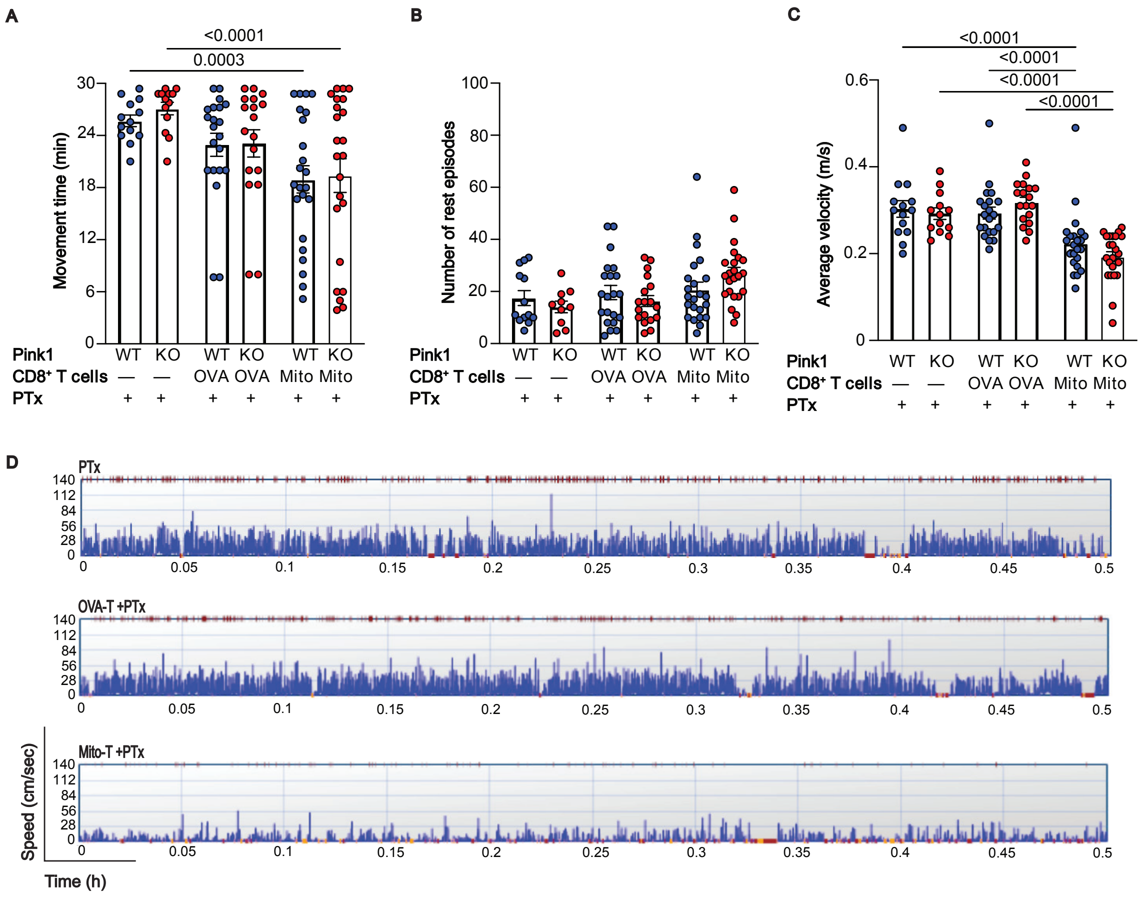


**Supplementary Figure 2**. **Motor impairment but not depressive-like behaviors after adoptive transfer of mitochondrial antigen-specific CD8$^{+}$ T. A.** Movement time (time of ambulation or stereotypic movement). **B.** Rest episode count (number of periods of inactivity greater than or equal to 3s). **C.** Peak average velocity of ambulatory episodes (the highest value of the mean velocities for each ambulatory episode). **D.** Sample velocity plot generated by Fusion from locomotor data, represent the velocity over period of 30 min that represents the peak average velocity. Red dots over the plot represent the time of vertical activity episode.

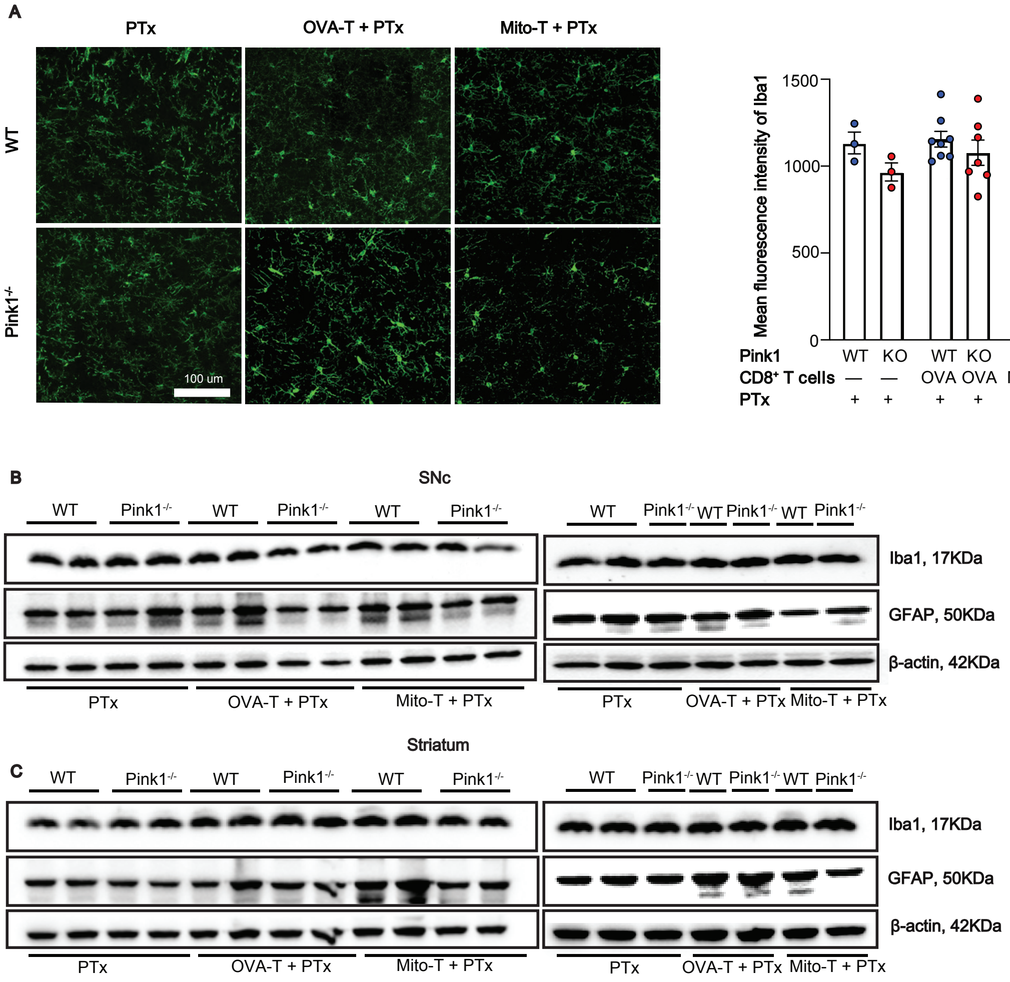


**Supplementary Figure 3. Lack of evidence for glial activation after mitochondrial antigen-specific CD8$^+$ T-cells adoptive transfer. A.** Representative immunohistochemistry image showing iba1+ cells in the dorsal striatum (left) and a compilation of the mean fluorescent intensity of iba1 (right). **B.** Western blot showing expression of iba1 and GFAP in the SNc. **C.** Western blot showing expression of iba1 and GFAP in the striatum. The housekeeping protein β-Actin was used loading control. The position and molecular weight of each protein is indicated.
